# MRN and Topoisomerase IIIα-RMI1/2 synchronize DNA resection motor proteins

**DOI:** 10.1101/2022.07.01.498452

**Authors:** Michael M. Soniat, Giaochau Nguyen, Hung-Che Kuo, Ilya J. Finkelstein

## Abstract

DNA resection—the nucleolytic processing of broken DNA ends—is the first step of homologous recombination. Resection is catalyzed by the resectosome, a multi-enzyme complex that includes BLM helicase, DNA2 or EXO1 nucleases, and additional DNA-binding proteins. Although the molecular players have been known for over a decade, how the individual proteins work together to regulate DNA resection remain unknown. Using single-molecule imaging, we characterized the roles of MRN and TOP3A-RMI1/2 during long-range DNA resection. BLM partners with TOP3A-RMI1/2 to form the BTRR complex (or BLM dissolvasome). TOP3A-RMI1/2 aids BLM in initiating DNA unwinding, and along with MRN, stimulates DNA2-mediated resection. Furthermore, MRN promotes the association between BTRR and DNA, and synchronizes BLM and DNA2 translocation to prevent BLM from pausing during resection. Together, this work provides direct observation of how MRN and DNA2 harness the BTRR complex to resect DNA efficiently and how TOP3A-RMI1/2 regulates BLM’s helicase activity to promote efficient DNA repair.

## Introduction

Homologous recombination (HR) is one of two major eukaryotic double-strand DNA break (DSB) repair pathways. HR uses the intact sister chromatid during the S/G2 phase to promote error-free repair of DSBs (1, 2). HR initiates when the resection machinery, termed the resectosome, assembles to process (resect) the genome to generate kilobase-length stretches of single-stranded DNA (ssDNA) (3–12). The resectosome is a multi-enzyme complex composed of a helicase, a nuclease, and regulatory proteins. In humans, resection is initiated when MRE11-RAD50-NBS1 (MRN) and CtIP make an initial incision at the DSB (11, 13). This aids in the assembly of the core resectosome consisting of the Bloom Syndrome Helicase (BLM) along with Exonuclease 1 (EXO1) or DNA2 nuclease/helicase (**Figure 1A**) (7, 10). The resulting ssDNA produced by the resectosome is rapidly bound by the ssDNA-binding protein Replication Protein A (RPA), which protects the ssDNA from degradation before being replaced by the RAD51 recombinase for downstream homology search (14–17).

**Figure 1.**
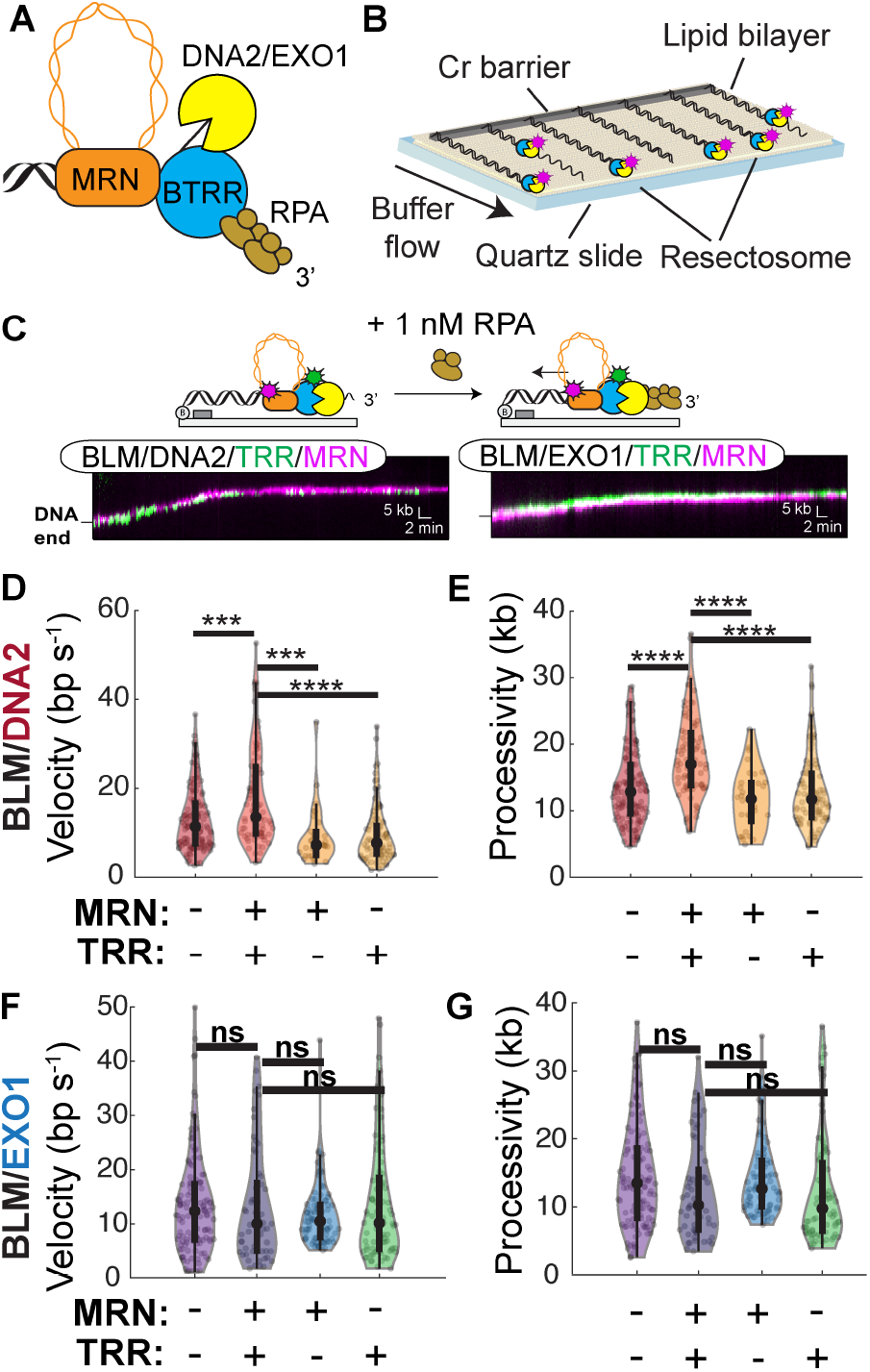
MRN and TOP3A-RMI1/2 stimulate DNA2-mediated resection. (A) Schematic of the human resectosome, consisting of MRE11-RAD50-NBS1 (MRN; orange), the nucleases DNA2 or EXO1 (yellow), BLM-TOP3A-RMI1/2 (blue), and Replication Protein A (RPA; brown). (B) Schematic of single-molecule resection assay (C) Representative kymograph of MRN (magenta), BTRR (green), and DNA2 or EXO1 resecting DNA. (D) Velocities and (E) processivities of BLM/DNA2 with and without TOP3A-RMI1/2 or MRN complex (N>50 for all experiments). Black bars show the interquartile range (thick bars) and 95% confidence intervals (thin bars). The black dot in the middle is the median. (F) Velocities and (G) processivities of BLM/EXO1 with and without TOP3A-RMI1/2 or MRN complex (N>50 for all experiments). (not significant; ns, p>0.05; ****, p<0.0001).

Accessory proteins regulate DNA resection to both initiate in a timely manner and to prevent over-resection and subsequent loss of genetic information (8, 12, 18). For example, in *S. cerevisiae*, the Sgs1 helicase (BLM homolog) and Dna2 nuclease/helicase interact with Mre11-Rad50-Xrs2 (MRX; the MRN homolog) and Top3-Rmi1 for efficient DNA resection (19, 20). The MRN complex has both exo- and endonuclease activity which is important for initial processing and the removal of protein adducts at DSBs (21–26). In addition to its nuclease-dependent roles, MRN/X also stimulates DNA resection via an incompletely understood nuclease-independent mechanism (19, 20, 27– 32). The human homologs of the *S. cerevisiae* Top3-Rmi1 are TOP3A (a type 1A topoisomerase) and the heterodimer RMI1-RMI2 (yeast don’t encode an RMI2 homolog). These proteins interact with BLM to form the BTRR complex (also known as the BLM dissolvasome) (33– 39). BTRR participates in DNA resection, double Holliday junction dissolution, resolution of ultra-fine bridges, replication fork reversal, and interstrand crosslink repair (40–43). During the initial stages of HR, the BTRR complex promotes long-range DNA2-mediated resection (44, 45). In later stages of HR, following RAD51 loading and strand invasion, BTRR is critical for the branch migration of Holliday junctions to form a hemicatenane, followed by decatenation by TOP3A-RMI1/2, resulting in non-crossovers (46, 47). Though the role of TOP3A-RMI1/2 is well defined in the later stages of HR, their precise roles in DNA resection remain unclear. For example, a catalytically inactive TOP3A mutant stimulates long-range DNA resection *in vitro*, suggesting that TOP3A-RMI1/2 may play a non-enzymatic role in resectosome assembly and/or translocation (20, 44).

Here, we use single-molecule fluorescence imaging to decipher the functions of individual resectosome components during DNA resection. MRN and TOP3A-RMI1/2 both help BLM to initiate DNA unwinding. MRN and TOP3A-RMI1/2 also stimulate DNA2-mediated resection. Finally, MRN synchronizes the translocation speeds of BLM and DNA2 to prevent BLM pausing. We reveal that MRN and TOP3A-RMI1/2 are regulatory resectosome components that initiate DNA resection and synchronize the individual motors during kilobase-long DNA processing.

## Results

### MRN and TOP3A-RMI1/2 together stimulate the DNA2 resectosome

To understand how MRN and TOP3A-RMI1/2 regulate DNA processing, we adapted our single-molecule resection assay to quantify the movement of DNA2 or EXO1 in complex with MRN and BLM-TOP3A-RMI1/2 (BTRR; **Figure 1**) (28, 48–50). We purified RMI1/2 with an N-terminal FLAG epitope on the RMI2 subunit. TOP3A-RMI1/2 was reconstituted by mixing TOP3A and RMI1/2 in a 1:3 ratio, followed by size exclusion chromatography. These three proteins formed a stable complex that eluted as a single peak on a Superose 6 column (**Figure S1**). TOP3A-RMI1/2 was then mixed with BLM to assemble the BTRR resectosome and labeled with fluorescent anti-FLAG antibodies (targeting RMI2 as described above). Biotinylated MRN was conjugated with streptavidin quantum dots (QDs) that emit in spectrally distinct channels (**Figure S1**). We have previously confirmed that fluorescent MRN retains its biochemical activities (28). For the single-molecule resection assay, 48.5 kb-long double-stranded DNAs with biotin on one end and a 78 nt 3’-overhang on the opposite end are organized on the surface of a microfluidic flowcell (**Figure 1B**)(49, 51). Fluorescent BTRR complex is incubated with MRN and DNA2 or EXO1 before being injected into flowcells for single-molecule imaging. As expected, MRN and BTRR in complex with DNA2 and EXO1 bound the free DNA ends and resected the DNA in the presence of 1 nM RPA (**Figure 1C**). The MRN/BTRR/DNA2 complex resected DNA for 18 ± 6 kb (mean ± st. dev; n=82) with a velocity of 18 ± 11 bp s^-1^. Omitting either TOP3A-RMI1/2 or MRN decreased BLM/DNA2 velocity ∼2-fold (BTRR/DNA2: 9 ± 6 bp s^-1^, n=94 molecules; MRN/BLM/DNA2: 9 ± 7 bp s^-1^, n=30) and decreased processivity by 1.4-fold (BTRR/DNA2: 13 ± 5 kb; MRN/BLM/DNA2: 12 ± 5 kb) (**Figure 1C-E and Supplementary Table 1**). Our results with the human resectosome are consistent with the stimulation of *S. cerevisiae* Sgs1-Dna2 by MRX and Top3-Rmi1 (19, 20). In contrast, the addition of MRN and TOP3A-RMI1/2 to BLM/EXO1 resectosomes did not change the processivity or velocity (∼13 ± 7 kb; ∼13 ± 9 bp s^-1^; n>50 for all conditions), suggesting that MRN and TOP3A-RMI1/2 selectively regulate DNA2-mediated resection (**Figure 1F**,**G and Supplementary Table 1**).

### MRN and TOP3A-RMI1/2 recruit DNA2 to free DNA ends

In addition to its nucleolytic activity, DNA2 also encodes a 5’→3’ helicase domain that can unwind kilobases of dsDNA (52–56). We first tested the importance of both DNA2’s nuclease and helicase activity in DNA resection with MRN and the BTRR complex on the 3’-overhang DNA substrate. As expected, the nuclease-deficient DNA2(D277A) inhibited DNA resection (processivity: 2 ± 2 kb; velocity: 2 ± 2 bp s^-1^, n=89) (**Figure 2A,B**). Furthermore, a helicase-deficient DNA2(K654R) mutant decreased resection processivity and velocity (processivity: 13 ± 6 kb; velocity: 11 ± 7 bp s^-1^, n=76). Omitting TOP3A-RMI1/2 and MRN abrogated the negative effects of DNA2(K654R) on resection processivity and velocity (14 ± 8 kb; 12 ± 8 bp s^-1^, n=42) (**Figure S2A**,**B**). These results show that both DNA2’s nuclease and helicase activity are required for rapid long-range DNA processing.

**Figure 2:**
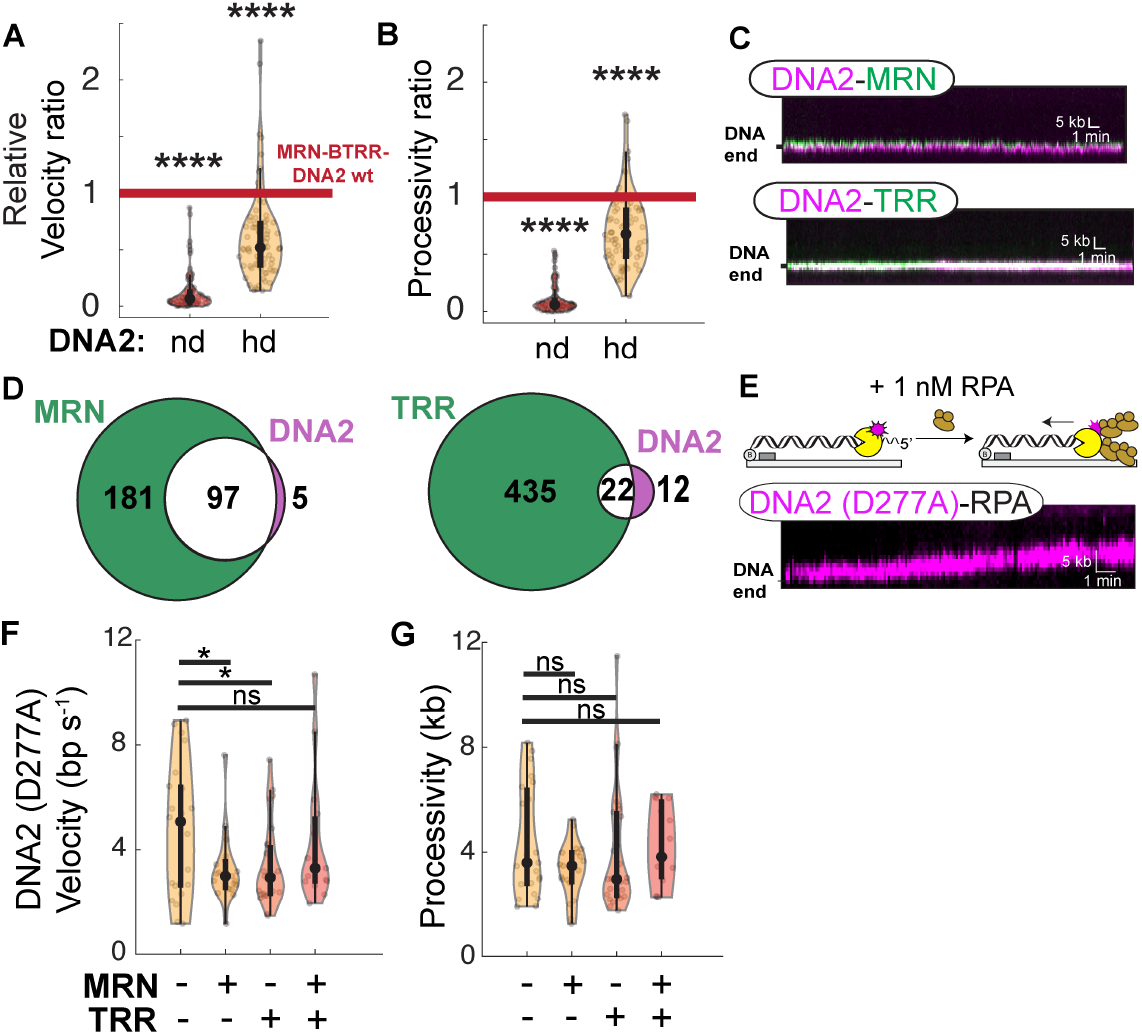
MRN and TOP3A-RMI1/2 recruits DNA2 to free DNA ends. (A) Ratio of MRN/BTRR/DNA2 velocities and (B) processivities with nuclease-deficient (nd) or helicase-deficient (hd) DNA2 mutants. Both velocity and processivity are compared to the ratios of MRN/BTRR/DNA2 wild-type complex from Figure 1D-E. (ns, p > 0.05; *p < 0.05; **p < 0.01; ***p < 0.001; ****p < 0.0001). (C) Representative kymographs of co-localization of DNA2 (magenta) with MRN and TOP3A-RMI1/2 (TRR; green) at DNA ends. MRN and TRR were imaged in different experiments to guarantee an unambiguous fluorescent signal. (D) Venn diagram that shows MRN and TOP3A- RMI1/2 co-localize DNA2 at free DNA ends. This interaction greatly increases the number of DNA-bound DNA2 molecules. (E) Representative kymograph showing helicase activity of nuclease deficient DNA2 (D277A) mutant in the presence of RPA. (F) Velocities and (G) processivities of DNA2 (D277A) helicase activity with and without TOP3A-RMI1/2 or MRN complex. (not significant; ns, p>0.05; *, p<0.05; **, p<0.01; ***, p<0.001; ****, p<0.0001).

Since TOP3A-RMI1/2 and MRN stimulate the BLM/DNA2 resectosome, we tested whether either TOP3A-RMI1/2 and/or MRN can stimulate DNA2 alone. MRN recruits DNA2 to DSBs in human cells but does not affect DNA2 nuclease activity *in vitro* (30). Consistent with this report, 95% of DNA2 molecules co-localized with MRN (n=97/102) at free DNA ends (**Figure 2C,D**). We saw a more modest 65% of TOP3A-RMI1/2 complexes (n=22/34) co-localizing with DNA2. DNA2 did not translocate beyond our spatial resolution of ∼500 bp with MRN and TOP3A-RMI1/2 (together or independently) in the presence of RPA. Previous studies showed that suppression of DNA2’s nuclease activity stimulates processive helicase activity on DNA substrates containing a 5’-flap in the presence of RPA (52–54). To observe the helicase activity, we monitored nuclease-deficient DNA2(D277A) on DNA substrates containing 12 nt 5’-overhang in the presence of 1 nM RPA, similar to previous studies (52, 54, 57). DNA2 was a processive helicase, motoring ∼4 ± 2 kb with a velocity of 5 ± 3 bp s^-1^ (n=23 molecules) (**Figure 2E-G and Supplementary Table 2**). The addition of MRN, TOP3A-RMI1/2, or both together had no effect on DNA2’s processivity. However, adding either MRN or TOP3A-RMI1/2 decreased DNA2’s velocity by ∼1.4-fold relative to DNA2(D277A) alone. In contrast, the addition of both MRN and TOP3A-RMI1/2 restored helicase activity to that of DNA2 alone. These results are broadly consistent with a model where MRN and TOP3A-RMI1/2 help DNA2 engage free DNA ends but do not stimulate its nuclease or motor activities.

### TOP3A-RMI1/2 helps BLM initiate DNA unwinding

Next, we investigated how TOP3A-RMI1/2 regulates BLM helicase. BLM was fluorescently labeled via a fluorescent anti-HA antibody directed to an N-terminal HA epitope, as we described previously (50). TOP3A-RMI1/2 co-localized with BLM at the free DNA ends (**Figure 3A**). We recently showed that RPA aids BLM in initiating helicase activity from 3’-ssDNA overhangs (50). When RPA is omitted, only ∼30% of BLM molecules initiate translocation. However, adding TOP3A-RMI1/2 increases the number of translocating BLM molecules ∼2.3-fold (77%; n=86/111) (**Figure 3B**). TOP3A was sufficient to recapitulate most of this stimulation (62%; n=83/133), whereas adding RMI1/2 alone slightly decreased the number of translocating BLM molecules (19%; n=34/182). Finally, adding RPA did not further stimulate the number of translocating BTRR complexes (83%; n=67/81). Surprisingly, although TOP3A-RMI1/2 initiated more BLM helicases, it also decreased BLM’s velocity ∼2-fold (14 ± 11 bp s^-1^; n=86) and slightly reduced processivity ∼1.2-fold (14 ± 8 kb) (**Figure 3C-D and Supplementary Table 3**). Furthermore, >50% of BTRR molecules dissociate from DNA during the 30-minute experiment (**Figure 4C**). BLM-TOP3A had a similar velocity and processivity as the BTRR complex (n=83). However, the processivity of BLM-RMI1/2 was further reduced ∼2-fold compared to the BTRR complex (8 ± 3 kb; n=34). Adding RPA did not change BTRR velocity but slightly decreased the average processivity (11 ± 6 kb; n=67). We conclude that TOP3A-RMI1/2 helps to initiate DNA unwinding but reduces BLM’s velocity and processivity.

**Figure 3:**
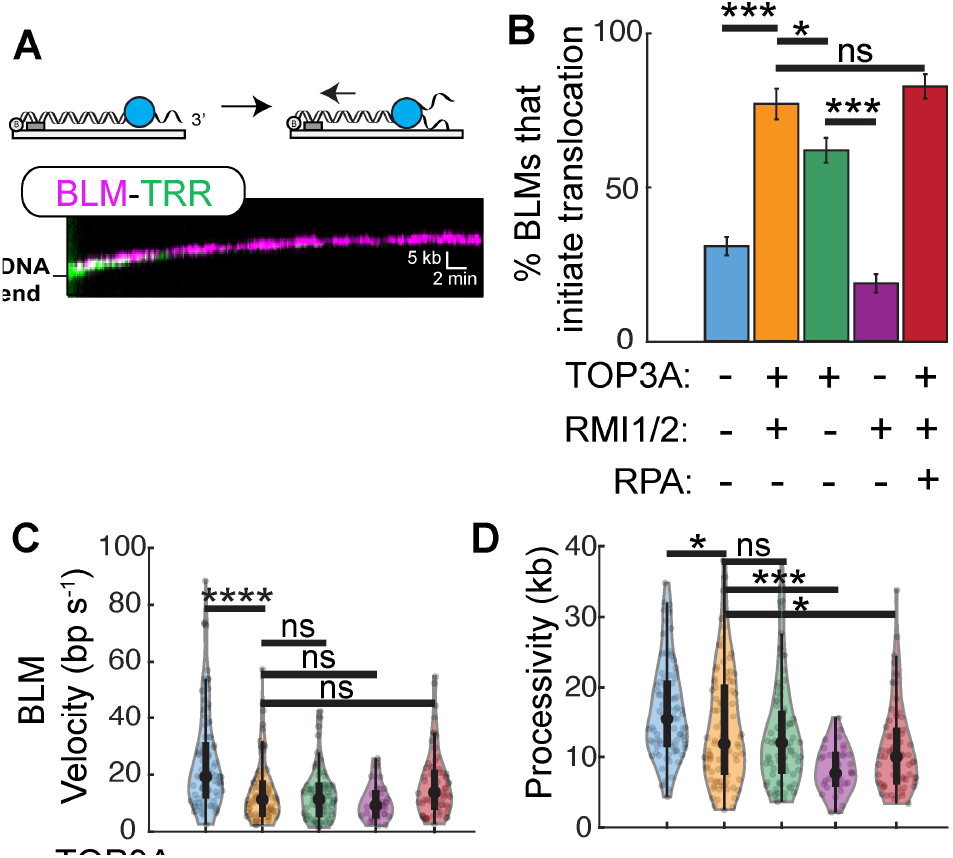
TOP3A-RMI1/2 complex promotes initiation of BLM helicase activity. (A) Representative kymographs showing helicase activity of BLM (magenta) with the TOP3A-RMI1/2 complex (green) along DNA. (B) TOP3A-RMI1/2 stimulates BLM helicase initiation. Error bars: S.D. as determined by bootstrap analysis. (C) Velocities and (D) processivities of the helicase activity of the BTRR complex. (not significant; ns, p>0.05; *, p<0.05; **, p<0.01; ***, p<0.001; ****, p<0.0001).

**Figure 4.**
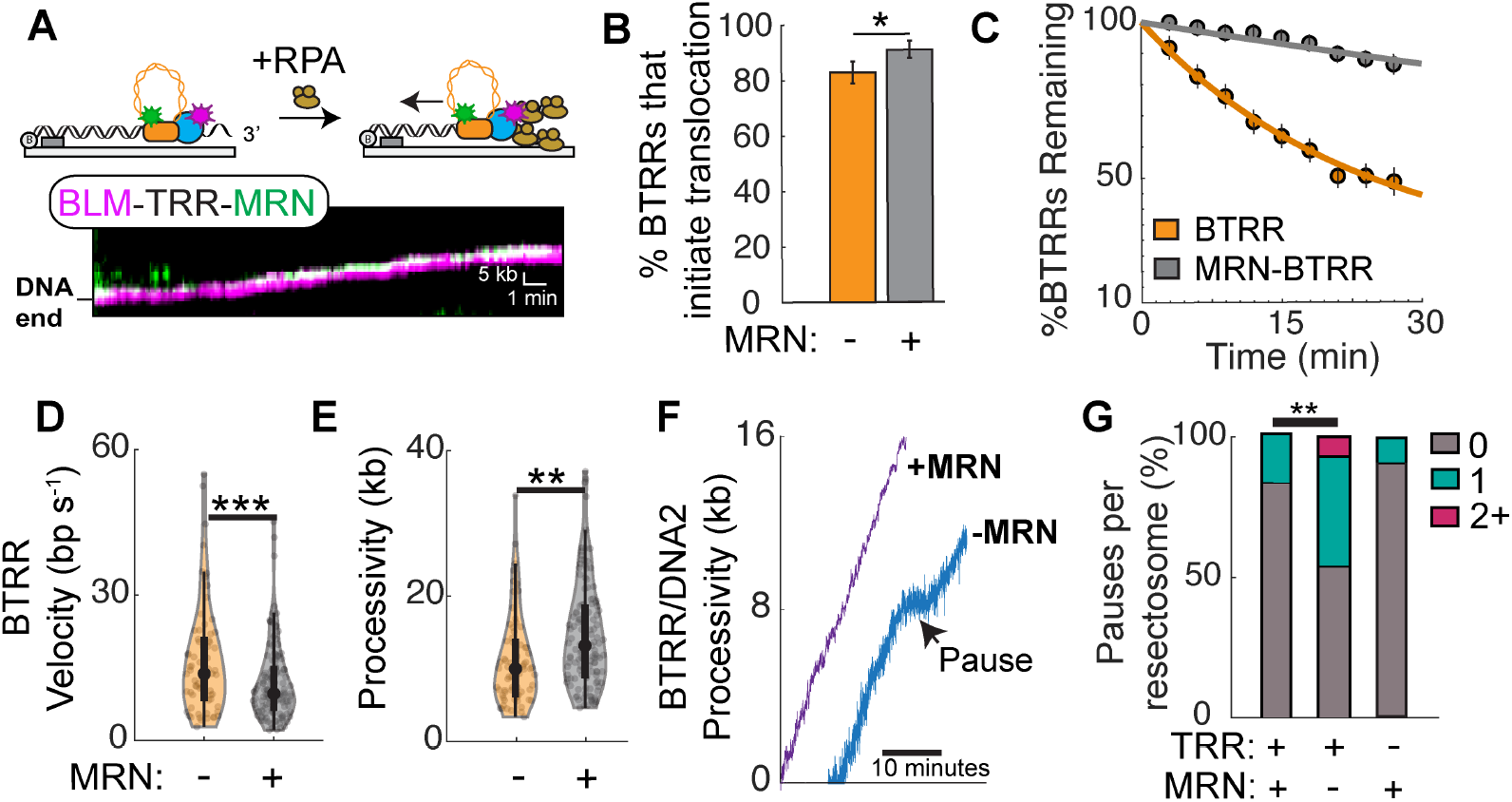
MRN prevents BTRR DNA dissociation. (A) Representative kymographs showing helicase activity of BLM (magenta) with the TOP3A- RMI1/2 complex (dark) and MRN (green) along DNA. (B) MRN has no additional effect on helicase initiation by the BTRR complex with RPA. Error bars: S.D. as determined by bootstrap analysis. (C) MRN retains the BTRR complex on DNA, as indicated by the lifetime analysis of the MRN-BTRR complex. In the presence of MRN, BTRR essentially doesn’t dissociate from the DNA during the 30 min experiment. (D) Velocities and (E) processivities of the BTRR complex helicase activity with or without MRN (F) Representative particle tracking of BTRR/DNA2 resection with (purple) and without MRN (blue). (G) MRN decreases pausing events during BTRR/DNA2 resection. (not significant; ns, p>0.05; *, p<0.05; **, p<0.01; ***, p<0.001; ****, p<0.0001).

### MRN prevents BTRR dissociation from DNA

Having established that TOP3A-RMI1/2 helps initiate DNA unwinding, we tested whether MRN further stimulates the BTRR complex. MRN is important for BLM recruitment to DSB and has been shown to stimulate unwinding activity (30, 58, 59). As expected, MRN co-localizes with BTRR and moves together with BTRR during DNA unwinding (**Figure 4A**). In the presence of 1 nM RPA, ∼90% (n=131/145) of MRN/BTRR resectosomes initiated DNA unwinding (**Figure 4B**). MRN also retained BLM molecules on the DNA (**Figure 4C**). In the presence of MRN, fewer than 30% of the BLM molecules dissociated from the DNA (n=131). This resulted in a 1.3-fold increase in processivity (15 ± 7 kb; n=131) but a ∼1.4-fold decrease in MRN/BTRR helicase velocity (12 ± 8 bp s^-1^; n=131) (**Figure 4D,E**). We conclude that MRN prevents the dissociation of the BTRR complex from DNA.

### MRN synchronizes the BLM and DNA2 motors

DNA2’s unwinding rate is ∼3-fold slower than BLM’s in the presence of RPA (**Figure 2F**). The yeast homolog of BLM, Sgs1, and Dna2 also show a ∼2-fold difference in unwinding rates (60). This difference in unwinding rates of BLM and DNA2 can lead to discoordination between the two motors. Consistent with this notion, we observed that ∼45% of the BTRR/DNA2 resectosomes paused for >30 seconds during DNA resection (n=42/94) with 12% (n=5/42) of these complexes pausing two or more times during their resection trajectories (**Figure 4G,H**). The change in resection velocity after the pause was heterogeneous and did not correlate with the pre-pause velocity (**Figure S2C**). Adding MRN suppressed these pauses (83% of resectosomes did not pause; n=68/82). MRN also suppresses pauses with the minimal MRN/BLM/DNA2 assembly (83% did not pause; n=25/30). We conclude that MRN coordinates BLM and DNA2 to stimulate efficient DNA resection.

## Discussion

DNA resection is catalyzed by either the EXO1 or DNA2 nucleases. Although EXO1 may be the predominant resection nuclease in human cells (48, 61–63), DNA2 is better at processing apurinic/apyrimidinic (AP) sites and 8-oxo-guanines (64). Furthermore, a super-resolution imaging study found comparable recruitment of both DNA2 and EXO1 at induced DSBs, suggesting that both nucleases are required during DNA resection (59). We had previously shown that MRN and BLM act as processivity factors for EXO1 in the presence of RPA (28, 50). We expand on the earlier study to show that TOP3A-RMI1/2 does not stimulate BLM/EXO1 resection. Instead, TOP3A-RMI1/2, in combination with MRN, stimulate DNA2-mediated resection. Additionally, MRN plays a scaffolding role by assembling the resectosome at a DSB and suppressing the dissociation of BTRR from DNA. MRN also prevents pausing by coordinating BLM and DNA2 during DNA resection **(Figure 5)**.

**Figure 5.**
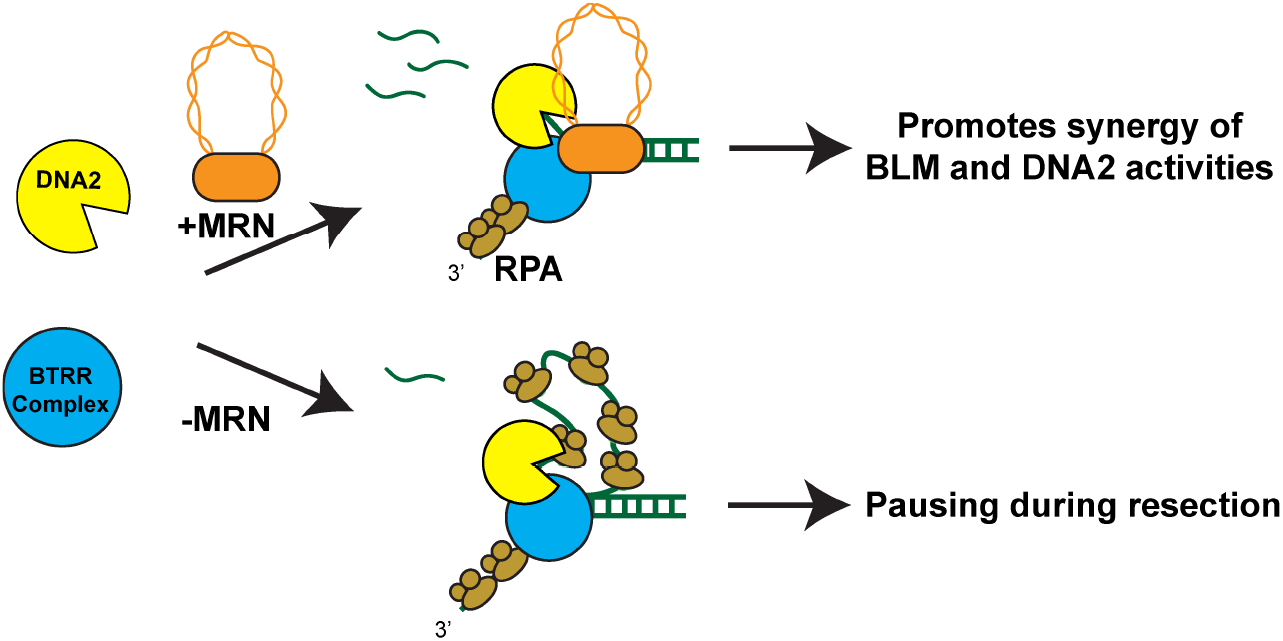
Model of MRN regulation of DNA resection motors.

Pausing during DNA resection has been observed with the bacterial RecBCD nuclease/helicase complex following recognition of the recombination hotspot sequence *χ* (crossover hotspot instigator-Chi) (65–67). RecBCD is functionally reminiscent of BTRR/DNA2 because it also encodes a fast (RecD) and slow (RecB) motor of opposite polarity. Two motors that move along opposite DNA strands with different speeds will generate a long ssDNA loop between them. Such loops are generated by the *E. coli* RecBCD helicase/nuclease due to differences in the translocation rates between the RecD and RecB motors (68). The molecular origin of RecBCD pausing stems from the slower motor “catching up” with the faster motor due to a conformational switch after *χ* recognition.

The underlying reason for pausing by BTRR/DNA2 is unknown, but it is unlikely to depend on a *χ*-like DNA sequence as sequence-dependent resection regulation has not been observed in yeast and human resectosomes. We conjecture that BTRR may unwind DNA in front of DNA2, leading to a growing ssDNA loop that ultimately pauses the entire complex. In this model, MRN prevents the accumulation of such ssDNA loops by synchronizing BLM and DNA2 helicase velocities. In support of this hypothesis, a recent single-molecule study showed that BLM retains contact with ssDNA as it unwinds double-stranded DNA (69). Additional high-resolution electron microscopy and other biochemical studies will be required to directly image an ssDNA loop between BLM and DNA2.

MRE11 and BLM co-localize early at DNA breaks immediately following damage in cells (59). This is consistent with our results showing that MRN assembles the resectosome at a DNA break. However, MRE11 and BLM do not associate as closely in the later stages of DNA resection (59). MRN’s role may thus be critical in the initiation and early coordination of BLM and DNA2. In the later stages of DNA resection, pausing may act as a negative DNA resection signal. Such pauses slow resection, possibly limiting over-resection and giving RAD51 sufficient time to complete the homology search. Together, this work shows that its conserved accessory factors regulate BLM’s helicase activity and that coordination with MRN and DNA2 stimulates DNA resection and, ultimately, efficient HR.

## Materials and Methods

### Protein Cloning and Purification

Oligonucleotides were purchased from IDT. Human RPA (pIF47) was purified from *E. coli* using a pET expression vector (48, 70, 71). Epitope-tagged human Exonuclease 1 (EXO1; pIF7) and MRN (pIF926) were purified from insect cells as previously described (28, 48, 50, 72, 73). FLAG-tagged human DNA2 nuclease-deficient (D277A) and helicase-deficient (K654R) mutants were generated by Phusion site-directed mutagenesis (Thermo Fisher) of wild-type DNA2-FLAG (pIF494) using oligos MS0015 and MS0016 for D277A (pIF495) and MS0017 and MS0018 for K564R (pIF496) (**Supplementary Table 4**). Wild-type DNA2, DNA2 (D277A), and DNA2 (K654R) were purified from insect cells as previously described (50, 73).

For single-molecule fluorescent imaging, 3xHA-BLM-His_6_ (pIF527) was expressed in Sf21 insect cells infected using the Bac-to-Bac expression system (Life Tech) (74). Cells were harvested 72 hours after infection, pelleted, frozen, and stored at -80°C. Cells were homogenized in buffer A containing 50 mM Tris-HCl pH 7.5, 500 mM NaCl, 10% glycerol, 2 mM β-mercaptoethanol, 10 mM imidazole, and 250 mM phenylmethane sulfonyl fluoride (PMSF) in a Dounce homogenizer (Kimble Chase; Kontes) followed by sonication on ice. Insoluble material was pelleted for 1 hr at 100,000 x g and the supernatant was added to Ni-NTA resin (QIAGEN, 30410) in batch and was eluted with an imidazole gradient containing 50 mM Tris-HCl pH 7.5, 500 mM NaCl, 10% glycerol, 2 mM β-mercaptoethanol, and 10-250 mM Imidazole. BLM fractions were then loaded on a 1 mL HiTrap Heparin (GE Healthcare) and eluted with a gradient from buffer B (50 mM Tris-HCl pH 7.5, 100 mM NaCl, 10% Glycerol, 1 mM DTT) to buffer C (50 mM Tris-HCl pH 7.5, 1 M NaCl, 10% Glycerol, 1 mM DTT). BLM was further purified using a Superose 6 (GE Healthcare) in buffer D (50 mM Tris-HCl pH 7.5, 200 mM NaCl, 10% Glycerol, 1 mM DTT).

pRSF-Duet plasmids containing His_6_-TOP3A and RMI1-His_6_/FLAG-RMI2 were kindly provided by Patrick Sung and Jim Daley. The TOP3A plasmid was transformed into Rosetta (DE3) pLysS cells and grown in LB media and induced with 0.2 mM IPTG for 18 hours at 16°C. Cells were pelleted and resuspended in buffer E (50 mM Tris-HCl pH 7.5, 1M KCl, 10% glycerol, and 1mM DTT) supplemented with 0.01% Igepal and 1mM PMSF. The cells were sonicated and the soluble material was clarified by centrifugation at 35,000 RCF for 45 minutes. The supernatant was added to Ni-NTA (QIAGEN, 30410) in batch, washed with buffer A supplemented with 20 mM imidazole, and eluted with buffer F (50 mM Tris-HCl pH 7.5, 200 mM KCl, 10% glycerol, 1mM DTT) supplemented with 250mM imidazole. TOP3A was further purified using a 1 mL HiTrap SP (GE Healthcare) with a gradient from buffer G (50 mM Tris-HCl pH 7.5, 50 mM KCl, 10% glycerol, 1mM DTT) to buffer H (50 mM Tris-HCl pH 7.5, 1 M KCl, 10% glycerol, 1mM DTT) and dialyzed overnight at 4°C in buffer I (50 mM Tris-HCl pH 7.5, 200 mM KCl, 10% glycerol, 1mM DTT).

The RMI1-His_6_/FLAG-RMI2 plasmid was transformed into Rosetta (DE3) pLysS cells and grown in LB media and induced with 0.2 mM IPTG for 18 hours at 16°C. The cells were sonicated, centrifuged, and purified by Ni-NTA resin similar to TOP3A. RMI1/2 elution from Ni-NTA purification were diluted to 50mM KCl with buffer J (50 mM Tris-HCl pH 7.5, 10% glycerol, 1 mM DTT), loaded on a 5mL HiTrap Q XL (GE Healthcare), and eluted with a gradient with buffers G and H. RMI1/2 was further purified using a Superdex 200 Increase (GE Healthcare) in buffer I. The TOP3A-RMI1/2 complex was assembled by incubating purified TOP3A and RMI1/2 (1:3 ratio) followed by purification using a Superdex 200 Increase (GE Healthcare) in buffer I.

### Single-Molecule Fluorescence Microscopy

All single-molecule data were collected on a Nikon Ti-E microscope in a prism-TIRF configuration equipped with a Prior H117 motorized stage. Flowcells were loaded into a custom-designed stage insert incorporating a chip mount, fluidic interface, and heating element (49). All experiments were maintained at 37°C by a combination of an objective heater (Bioptechs) and a custom-built stage-mounted heating block. The flowcell was illuminated with a 488 nm laser (Coherent) through a quartz prism (Tower Optical Co.). Data were collected with a 200 ms exposure, 2-second shutter (Vincent Associates) resulting in 1,800 frames in 1 hour, through a 60X water-immersion objective (1.2NA, Nikon), a 500 nm long-pass (Chroma), and a 638 nm dichroic beam splitter (Chroma), which allowed two-channel detection through two EMCCD cameras (Andor iXon DU897, cooled to -80°C). Images were collected using NIS-Elements software and saved in an uncompressed TIFF file format for later analysis (see below).

Fluorescent particles were tracked in ImageJ using a custom-written particle tracking script (available upon request). The resulting trajectories were analyzed in Matlab (Mathworks). Trajectories were used to calculate velocity and processivity for resectosomes. DNA substrates for single-molecule studies contained a 78 nt 3’-overhang or 12 nt 5’-overhang. These were prepared by annealing oligonucleotides IF007 and LM003 (3’-overhang) or IF007 only (5’-overhang) (**Supplementary Table 4**) (28).

#### Fluorescent protein labeling

3xHA-BLM (40 nM) were conjugated to Quantum Dots (QDs) pre-incubated with a rabbit anti-HA antibody (ICL Lab) on ice for 10 minutes in 20 μL. Next, BLM was incubated with the anti-HA QDs at a ratio of 1:2 for an additional 10 minutes on ice, diluted with imaging buffer (40 mM Tris pH 8.0, 60 mM NaCl, 200 µg/mL BSA, 2 mM DTT, 2 mM MgCl_2_, 1 mM ATP) to 200 µL and injected into the flowcell. FLAG- TOP3A-RMI1/2 or FLAG-DNA2 were labeled with QDs pre-incubated with a mouse anti-FLAG antibody (Sigma-Aldrich) on ice for 10 minutes prior to injection. Additionally, biotin-MRN was labeled via streptavidin QDs. Saturating biotin was added to the protein-QD conjugates to bind free streptavidin sites prior to injection

### Quantification and Statistical Analysis

For Figures 1-4, n represents the number of molecules. Quantification and statistical analyses were done using MATLAB (version: R2015b). Fluorescent particles were tracked using an in-house ImageJ script (available upon request) where the positions of individual molecules on DNA were determined by fitting the point spread function to a 2D Gaussian. Trajectories were used to calculate the velocity and processivity for BLM and DNA resection complexes. Statistical details of experiments can be found in the Results and figure legends where indicated.

## Data and Software Availability

All custom MATLAB and FIJI scripts are available upon request.

## Contact for Reagent and Resource Sharing

Further information and requests for resources and reagents should be directed to and will be fulfilled by the Lead Contact, Ilya Finkelstein

## Acknowledgments

We thank Tanya Paull, Patrick Sung, Jim Daley, and Marc Wold for reagents. We also thank members of the Finkelstein lab for comments on the manuscript

## Author Contributions

M.M.S. and G.N. prepared proteins and DNA. M.M.S. conducted all single-molecule experiments. M.M.S. and H.C.K. performed data analysis. M.M.S and I.J.F. secured funding, directed the project, and co-wrote the paper with input from all co-authors.

## Funding information

This work was supported by the NIH (GM120554 to I.J.F, CA092584 to I.J.F), CPRIT (R1214 to I.J.F.), the Welch Foundation (F-l808 to I.J.F.), and the American Cancer Society (PF-17-169-01-DMC to M.M.S.). The content is solely the responsibility of the authors and does not necessarily represent the official views of the National Institutes of Health.

## Conflicts of interest

The authors declare no competing interests.

## Notes

### Competing Interest Statement

The authors have declared no competing interest.

