## Supplemental Material for "MRN and Topoisomerase IIIα-RMI1/2 synchronize DNA resection motor proteins"

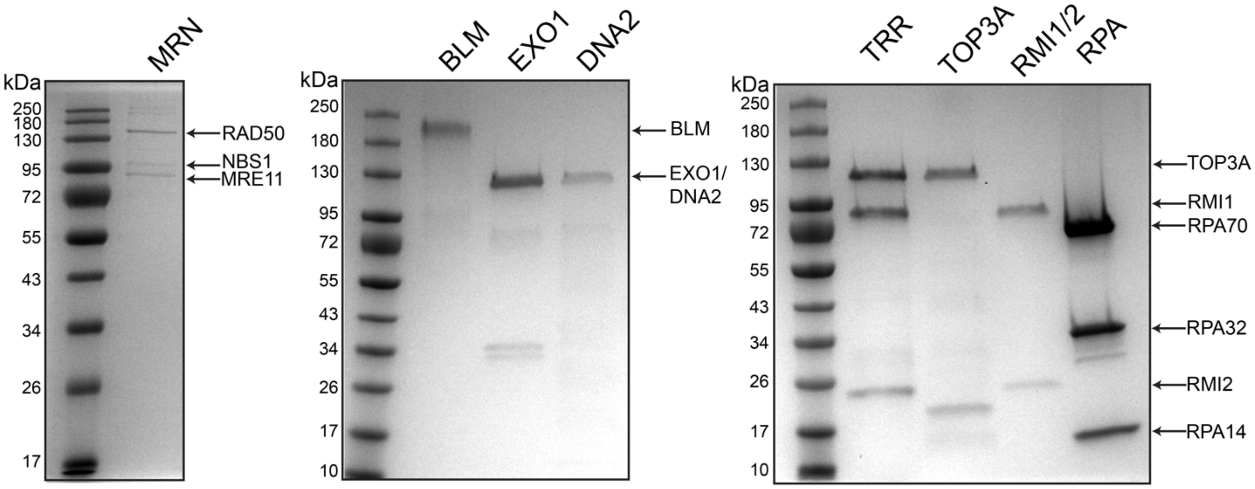


**Supplemental Figure 1:** SDS-PAGE gels of the recombinant proteins used in this study.

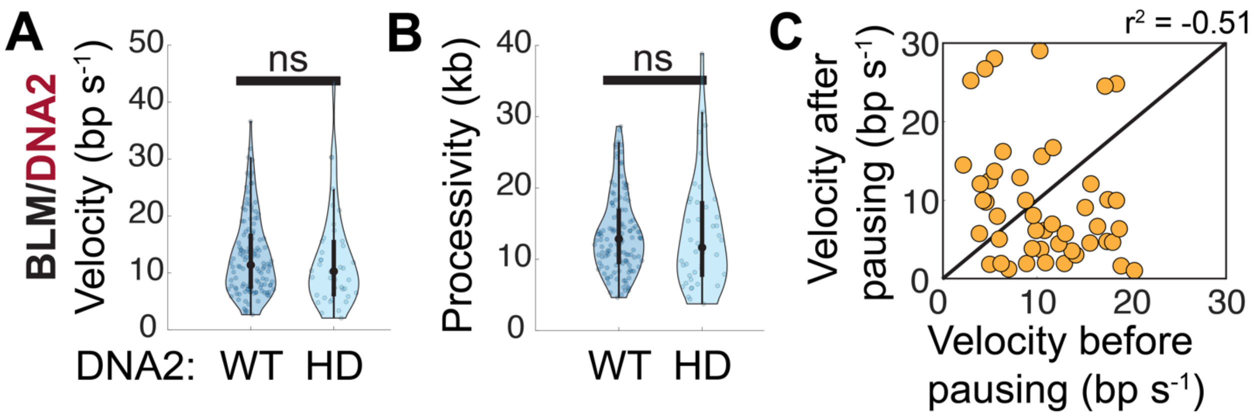


**Supplemental Figure 2: (A)** Velocities and **(B)** processivities of the BLM/DNA2 WT and the helicase-deficient DNA2(K654R) mutant (HD). Black bars show the interquartile range (thick bars) and 95% confidence intervals (thin bars). The black dot in the middle is the median (not significant; ns). **(C)** Velocities of individual BTRR/DNA2 complexes before and after pausing. Dashed line is shown as a reference with a slope of m = 1 (n = 42 complexes).

**Table S1**: **Velocity and processivity for DNA2 and EXO1-mediated DNA resection**

| Sample | Processivity (kb) mean ± st. dev | Velocity (bp $s^{-1}$) mean ± st. dev | Number of molecules (n) |
|---|---|---|---|
| BLM/DNA2 | 13±6 | 13±7 | 126 |
| MRN/BLM/DNA2 | 12±5 | 9±6 | 30 |
| BTRR/DNA2 | 13±5 | 9±6 | 94 |
| MRN/BTRR/DNA2 | 18±6 | 18±11 | 82 |
| BLM/EXO1 | 15±7 | 13±9 | 124 |
| MRN/BLM/EXO1 | 14±6 | 12±7 | 82 |
| BTRR/EXO1 | 12±8 | 14±11 | 79 |
| MRN/BTRR/EXO1 | 12±7 | 13±10 | 57 |
| MRN/BTRR/DNA2 (D277A) | 2±2 | 2±2 | 89 |
| MRN/BTRR/DNA2 (K654R) | 13±6 | 11±7 | 76 |
| BLM/DNA2 (K654R) | 14±8 | 12±8 | 42 |

**Table S2**: **Velocity and processivity for DNA2 helicase activity**

| Sample | Processivity (kb) mean ± st. dev | Velocity (bp $s^{-1}$) mean ± st. dev | Number of molecules (n) |
|---|---|---|---|
| DNA2 (D277A) | 4±2 | 5±3 | 23 |
| MRN/DNA2 (D277A) | 3±1 | 3±1 | 19 |
| TRR/DNA2 (D277A) | 4±2 | 3±2 | 27 |
| MRN/TRR/ DNA2 (D277A) | 4±1 | 4±2 | 15 |

**Table S3**: **Velocity and processivity for BLM helicase activity**

| Sample | Processivity (kb) mean ± st. dev | Velocity (bp $s^{-1}$) mean ± st. dev | Number of molecules (n) |
|---|---|---|---|
| BLM | 17±7 | 25±18 | 90 |
| BTRR | 14±8 | 14±11 | 86 |
| BLM/TOP3A | 13±8 | 13±9 | 83 |
| BLM/RMI1/2 | 8±3 | 10±6 | 34 |
| BTRR/RPA | 11±6 | 17±11 | 67 |
| MRN/BTRR/RPA | 15±7 | 12±8 | 131 |

**Table S4**: **Oligonucleotides used in this study.**

| Name | Sequence |
|---|---|
| IF007 | [p]AGG TCG CCG CCC[Bio] |
| LM003 | [p]GGG CGG CGA CCT TTTTTTTTTTTTTTTTTTTTTTTTTTTTTTTTTTTTTTTTTTTTTTTTT TTTTTTTTTTTTTTTTTTTTTTTTTTTTT |
| MS0015 | TATTTTGCCTTTCAATCCAAACCTAGG |
| MS0016 | GCCGTTACAGTTGGTGTGAAAATACATCGAG |
| MS0017 | TCCTGTCCCAGGCATACCC |
| MS0018 | CGTACAACTACGATATGTACTCTCGTAAGAATTC |
